# Membrane Orientation and Oligomerization of the Melanocortin Receptor Accessory Protein 2

**DOI:** 10.1101/2020.08.03.235200

**Authors:** Valerie Chen, Antonio E. Bruno, Laura L. Britt, Ciria C. Hernandez, Luis E. Gimenez, Alys Peisley, Roger D. Cone, Glenn L. Millhauser

## Abstract

The melanocortin receptor accessory protein 2 (MRAP2) plays a pivotal role in the regulation of several G-protein coupled receptors (GPCR) that are essential for energy balance and food intake. MRAP2 loss-of-function results in obesity in mammals. MRAP2 and its homolog MRAP1 have an unusual membrane topology and are the only known eukaryotic proteins that thread into the membrane in both orientations. In this study, we demonstrate that the conserved polybasic motif that dictates the membrane topology and dimerization of MRAP1 does not control the membrane orientation and dimerization of MRAP2. We also show that MRAP2 dimerizes through its transmembrane domain and can form higher order oligomers that arrange MRAP2 monomers in a parallel orientation. Investigating the molecular details of MRAP2 structure is essential for understanding the mechanism by which it regulates GPCRs and will aid in elucidating the pathways involved in metabolic dysfunction.

## INTRODUCTION

The melanocortin receptor accessory protein 2 (MRAP2) regulates several G-protein coupled receptors (GPCRs) that play critical roles in the regulation of energy homeostasis, and heterozygous MRAP2 variants have been identified in obese humans^1–4^. MRAP2 modulates the signaling of the melanocortin-4 receptor (MC4R), one of the five GPCRs in the melanocortin receptor family^1,5^. MC4R and MRAP2 are expressed in the paraventricular nucleus of the hypothalamus (PVN), a primary region for the control of food intake. MC4R is essential for energy homeostasis and heterozygous mutations in MC4R are the most common monogenic cause of human obesity^6,7^. In zebrafish, MRAP2 allows for developmental control of MC4R signaling^5^. Zebrafish MRAP2a, which is restricted to larval development suppresses MC4R signaling while MRAP2b, which is expressed in adult zebrafish, increases MC4R’s sensitivity to its agonist *α*-melanocyte-stimulating hormone. Furthermore, MRAP2 enhances signaling through MC4R *in vitro* and overexpression of MRAP2 in MC4R-containing PVN neurons leads to a reduction in food intake and increased energy expenditure in female mice^1,8^. Targeted deletion of MRAP2 in mice results in an obese phenotype, however, mice lacking only MC4R are more obese than mice lacking both MRAP2 and MC4R^1^. This suggests that there are other mechanisms by which MRAP2 also promotes feeding. It is now understood that MRAP2’s regulation over GPCRs is not limited to the melanocortin receptor family. MRAP2 has been shown to promote feeding through inhibition of the prokineticin receptor-1 as well as to decrease food intake through inhibition of the orexin receptor^9,10^. Additionally, MRAP2 regulates hunger sensing by potentiating ghrelin signaling through its interaction with the growth hormone secretagogue receptor 1a (GHSR1a) in the arcuate nucleus of the hypothalamus (ARC)^11,12^.

MRAP2, as well as its well-studied homolog MRAP1, are single-pass transmembrane proteins that can insert into the membrane in both orientations – N-terminal domain out or in (Figure 1A, 1B) ^13–15^. Like MRAP1, MRAP2 can homodimerize and form anti-parallel dimers^16,17^. MRAP2 and MRAP1 are the only two proteins in the eukaryotic proteome that are currently known to exhibit this unusual membrane orientation. The orientation of most membrane proteins is predicted by the “positive inside rule,” where the charged amino acids flanking the transmembrane domain determine the overall orientation such that the more positive region faces the cytosol ^18,19^. Based on this rule, MRAP1 is predicted to insert into the membrane in both orientations, which agrees with experimental data. Using the same line of logic, MRAP2 is predicted to have its N-terminal domain in the cytosol. Nevertheless, recent findings show that MRAP2 has dual topology. It is evident that the “positive inside rule” is not sufficient for explaining MRAP2’s membrane orientation.

**Figure 1.**
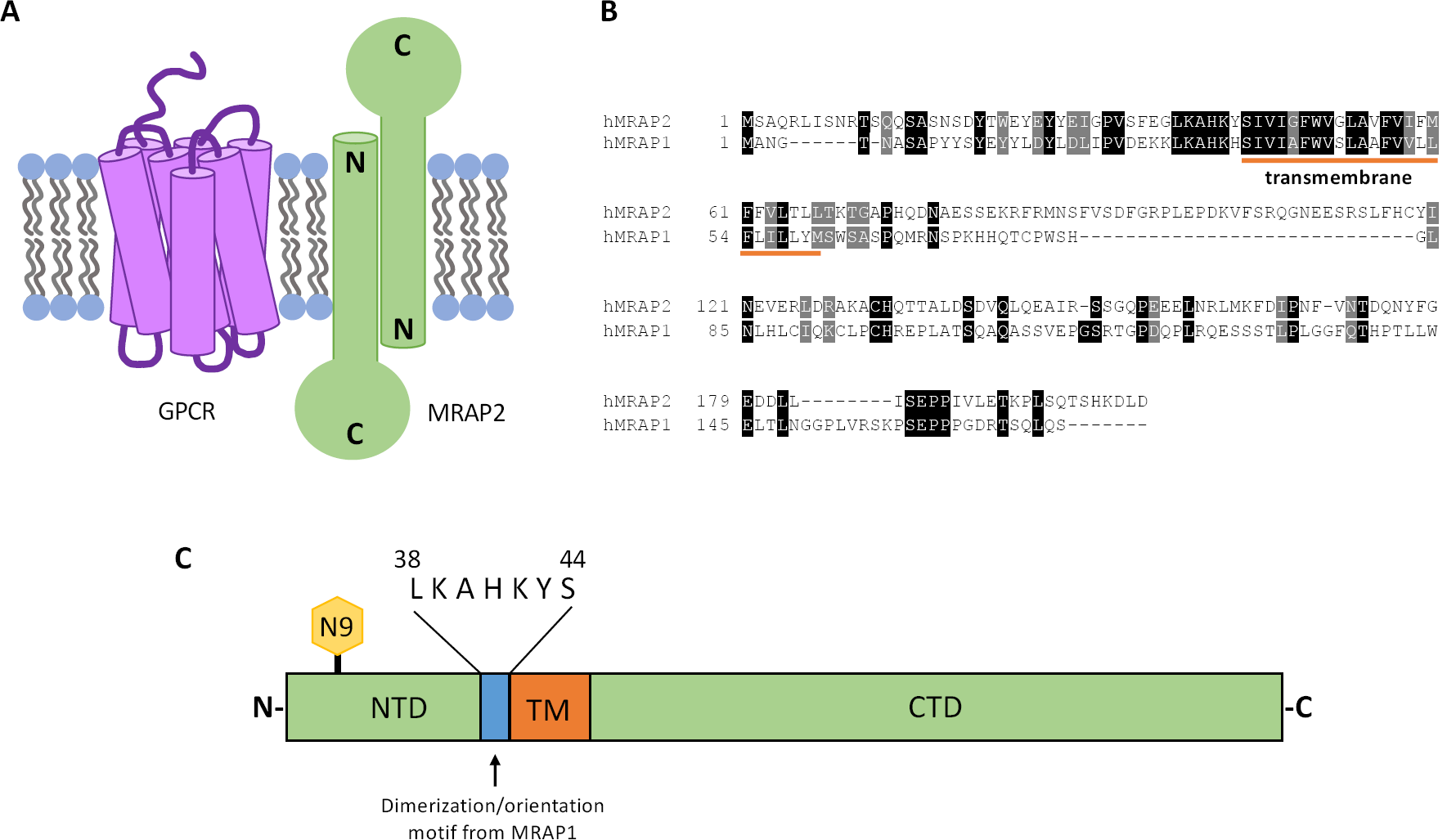
MRAP2 is an oligomeric, single-pass transmembrane protein that is critical for the modulation of GPCRs that are essential for energy homeostasis. A, Schematic depicting MRAP2’s dual topology. B, Protein alignment of human MRAP2 and human MRAP1. The transmembrane domain is underlined in orange. MRAP2 and MRAP1 are most conserved in their N-terminal and transmembrane domains. C, Schematic of MRAP2 protein. MRAP2 is glycosylated at residue 9, shown in yellow. The N-terminal domain (NTD) and C-terminal domain (CTD) are in green. The transmembrane (TM) domain is in orange. The conserved, polybasic motif required for the dual topology and dimerization of MRAP1 is in blue.

Despite the important role MRAP2 plays in the modulation of energy homeostasis, the sequence features within MRAP2 that dictate membrane orientation and dimerization are unknown. MRAP1’s membrane orientation and oligomeric state are dependent on a short polybasic segment adjacent to the transmembrane domain ^13,14^. While this motif is conserved in MRAP2, we show that this sequence does not dictate MRAP2’s membrane orientation or dimerization in cell culture. Additionally, using truncation mutations, we identify the transmembrane domain of MRAP2 as the minimal dimerization domain. Finally, we show that contrary to the assumption that MRAP2 can only form anti-parallel dimers, MRAP2 can form parallel dimers as well as higher order oligomers. Our results not only highlight important differences between MRAP1 and MRAP2 but also offer new insight into MRAP2 structure. Understanding the molecular details that determine MRAP2’s oligomeric state and membrane orientation will aid in elucidating the mechanism by which MRAP2 regulates GPCRs that are essential for metabolic processes.

## RESULTS

### The conserved polybasic motif (38-44) that is required for dual topology and dimerization of MRAP1 is not required for dual topology and dimerization of MRAP2

The dual topology and oligomeric state of MRAP1 is dictated by a short polybasic segment in the N-terminal domain that is directly adjacent to the transmembrane domain (Figure 1C). Sebag and Hinkle showed that mouse MRAP1 lacking residues 31 to 37 (MRAP1 Δ31-37) has a fixed membrane orientation such that the N-terminus of MRAP1 is extracellular and the C-terminus is cytosolic.^13^ This polybasic motif that determines the membrane orientation of MRAP1 is conserved in MRAP2. In order to investigate whether this sequence is required for dual topology of MRAP2, immunocytochemistry and microscopy were used to detect cell surface MRAP2 from intact, non-permeabilized cells that are transiently transfected with either N-terminally FLAG epitope-tagged or C-terminally FLAG epitope-tagged MRAP2 in human embryonic kidney 293T (HEK293T) cells (Figure 2A). Both the N-terminal domain and the C-terminal domain of wild-type MRAP2 were detected extracellularly. Unexpectedly, both the N-terminal domain and the C-terminal domain of MRAP2 lacking the polybasic motif (MRAP2 Δ38-44) were also detected on the cell surface. A control experiment was also performed to ensure formaldehyde fixation does not result in significant permeabilization of unpermeabilized cells (Figure S1). Immunocytochemistry and flow cytometry were used to measure the cell surface N-terminal domain to C-terminal domain ratio of wild-type MRAP2 Δ38-44 (Figure 2B). As expected, we find that wild-type MRAP2 has a cell surface N-terminal domain to C-terminal domain ratio of approximately 1. As a control, RAMP3, a single-pass transmembrane protein that was previously shown to favor a conformation with an extracellular N-terminus^13,20^ does in fact have an N-terminal domain to C-terminal domain ratio of approximately 2. We confirm that while deletion of the polybasic motif in MRAP2 does result in a modest preference for an extracellular N-terminal domain, MRAP2 Δ38-44 still has dual topology. Additionally, the presence of an immunoreactive “doublet band” for MRAP2 Δ38-44 indicates that the N-terminal asparagine residue exists in both glycosylated and unglycosylated forms (Figure 2C). This further indicates dual topology of MRAP2 Δ38-44 since the N-terminus must be in the ER lumen for glycosylation to occur. These results show that the conserved polybasic segment that is required for the dual topology of MRAP1 is not required for the dual topology of MRAP2.

**Figure 2.**
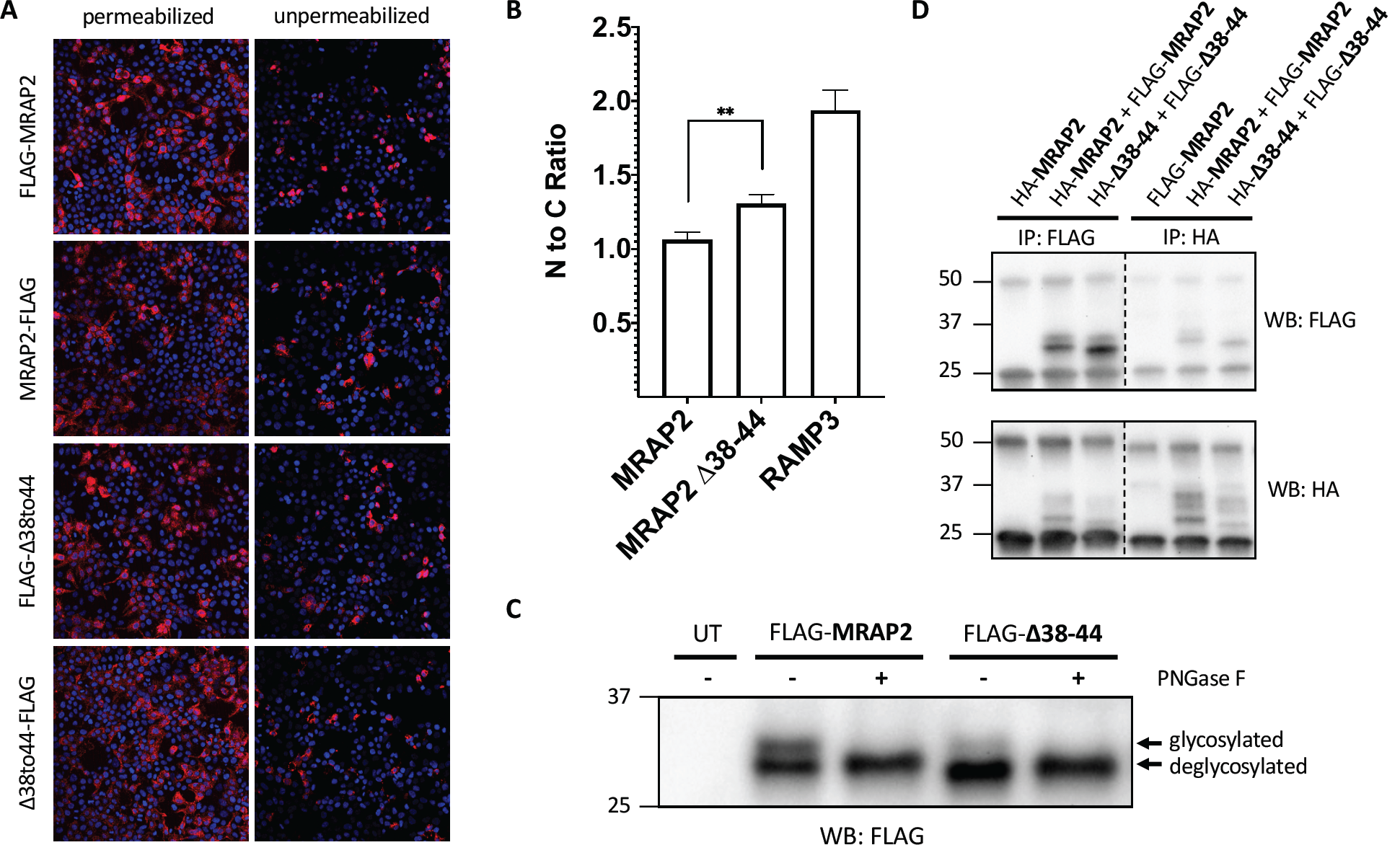
The conserved motif required for dual topology and dimerization of MRAP1 is not required for dual topology and dimerization of MRAP2. A, Both the N-terminus and C-terminus of MRAP2 WT and Δ38-44 are detected from intact, unpermeabilized HEK293 cells, as seen by immunofluorescence. FLAG-tagged MRAP2 is shown in pink and the nucleus is shown in blue. B, Flow cytometry was used to determine the N-terminal to C-terminal fluorescence ratio for MRAP2 WT, Δ38-44, and RAMP3 from intact cells expressing N-terminally tagged constructs and C-terminally tagged constructs. Expression levels for each construct were normalized using parallel experiments with permeabilized cells. Bars represent the mean from three independent experiments. Error bars are SEM. Statistical significance of differences was analyzed by *t* test. **p < 0.01 versus MRAP2. C, Immunoblot showing two protein species for FLAG-MRAP2 and FLAG-Δ38-44. Treatment with PNGase F results in a single, deglycosylated protein species. D, Immunoblot showing co-immunoprecipitation of HA- and FLAG-tagged MRAP2 and HA- and FLAG-tagged Δ38-44.

MRAP1 Δ31-37 cannot form dimers, likely due to the fact that it has a fixed membrane orientation.^13^ To determine whether MRAP2 Δ38-44 can form dimers or higher order oligomers, HEK293T cells were co-transfected with both FLAG- and HA-epitope tagged versions of either wild-type MRAP2 or MRAP2 Δ38-44. Co-immunoprecipitation, followed by western blotting showed that both wild-type MRAP2 and MRAP2 Δ38-44 form dimers or higher order oligomers (Figure 2D). The same experiment was performed in Chinese hamster ovary (CHO) cells and yielded the same results (Figure S2). Overall, these results indicate that the conserved polybasic motif that is required for dual topology and dimerization in MRAP1 is not required for either dual topology or dimerization of MRAP2.

### MRAP2 dimerizes through its transmembrane domain

MRAP2 is known to form dimers or higher order oligomers but the dimerization domain has not been identified. In order to identify the dimerization domain, constructs were created that either truncate the C-terminal domain (NTD-TM), the N-terminal domain (TM-CTD), or both the N- and C-terminal domains, leaving just the transmembrane domain (TM). HEK293T cells were co-transfected with both FLAG- and HA-epitope tagged constructs in the following combinations: TM-CTD + TM-CTD, NTD-TM + NTD-TM, NTD-TM + TM-CTD, and TM + TM. Co-immunoprecipitation, followed by western blotting show that neither the N-terminal domain nor the C-terminal domain are required for dimerization (Figure 3A). Specifically, TM-CTD can be co-immunoprecipitated with itself and with NTD-TM (Figure 3A, lanes 4, 6 and 7) and NTD-TM can be co-immunoprecipitated with itself (Figure 3A, lane 5). Finally, the transmembrane domain can also be co-immunoprecipitated with itself (Figure 3A, lane 8). The experiment was also performed in CHO cells and yielded the same results (Figure S2). Based on these results MRAP2 dimerizes through its transmembrane domain.

**Figure 3.**
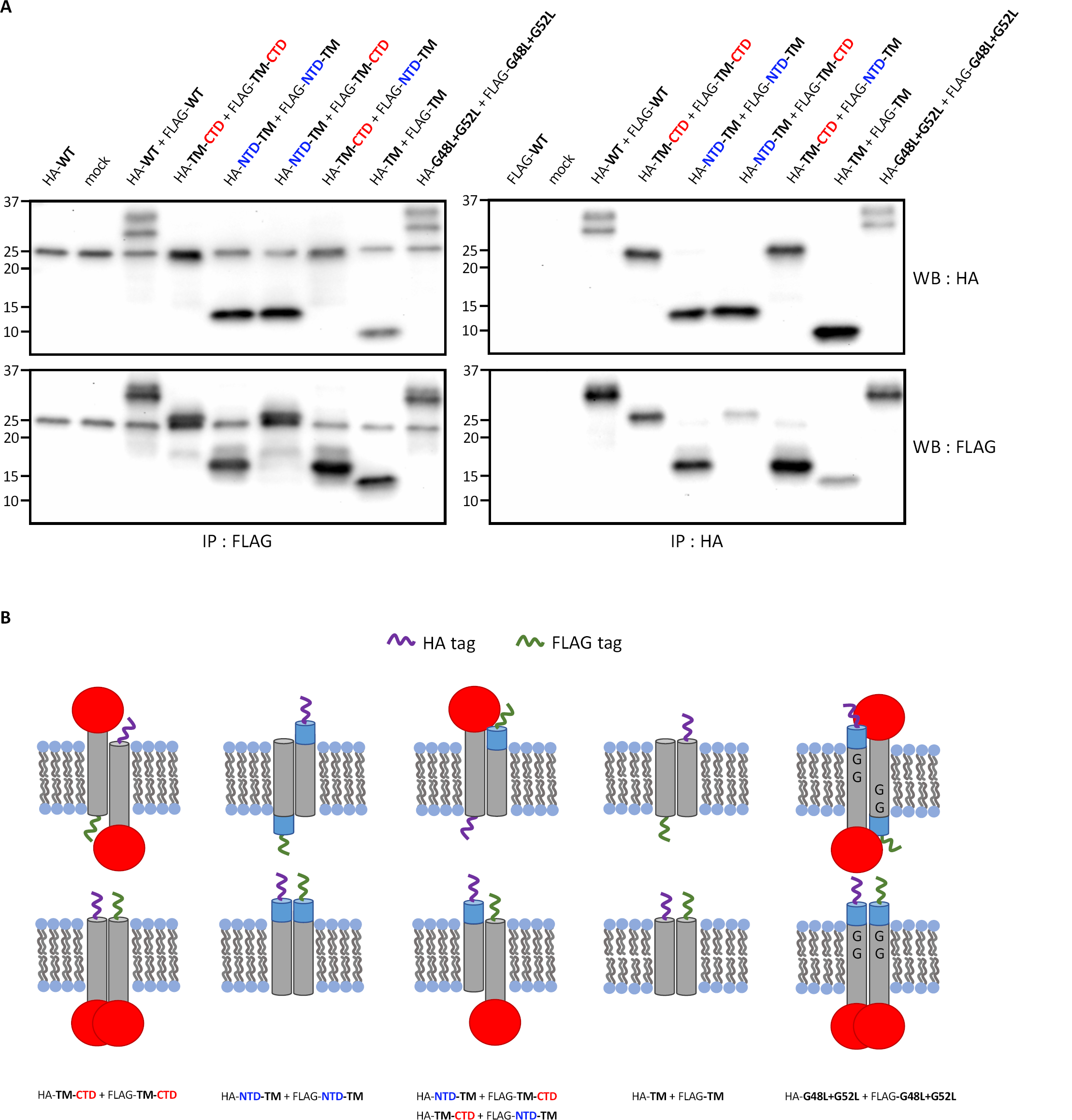
MRAP2 dimerizes through its transmembrane domain. A, HEK293 cells are co-transfected with HA- and FLAG-tagged versions of the following MRAP2 constructs: N-terminal domain truncation (TM-CTD), C-terminal domain truncation (NTD-TM), the transmembrane domain alone (TM), or a mutant that replaces the glycine residues within the transmembrane domain with leucine residues (G48L+G52L). Co-immunoprecipitations from cell lysates followed by immunoblotting show that neither the N-terminal domain, the C-terminal domain, nor the glycine residues in the transmembrane domain are required for dimerization of MRAP2. B, Schematic depicting the co-immunoprecipitated HA- and FLAG-tagged MRAP2 dimers.

The GxxxG motif is common in transmembrane helix interactions^21,22^. We also investigated whether the glycine residues within the transmembrane domain of MRAP2 are required for dimerization by mutating the glycine residues that make up this motif to leucine residues (G48L + G52L). Co-immunoprecipitations from cell lysates followed by immunoblotting also show that the glycine residues within the transmembrane domain are not required for MRAP2 dimerization (Figure 3A, lane 9).

### MRAP2 can form parallel dimers and higher order oligomers

Based on bimolecular fluorescence complementation experiments by Sebag and Hinkle, MRAP2, like MRAP1, forms anti-parallel dimers^16^. While previous experiments show that MRAP1 forms exclusively anti-parallel dimers and does not form parallel dimers, it is unclear as to whether this holds true for MRAP2. To determine whether MRAP2 forms parallel dimers, a NanoBiT protein-protein interaction assay was performed. The NanoBiT system is composed of a large BiT (LgBiT) and small BiT (SmBiT) that have very little to no luciferase activity on their own. However, when LgBiT and SmBiT are in close proximity within the cell, the functional luciferase will then generate a luminescent signal at even low protein expression levels, driven by weak HSV-TK promoter. The low intrinsic affinity between the isolated SmBiT and LgBiT lessens the likelihood of spurious dimerization driven by these segments. The following combinations of DNA were transfected into HEK293T cells: MRAP2-LgBiT + SmBiT-PRKACA, LgBiT-MRAP2 + SmBiT-PRKACA, LgBiT-MRAP2 + MRAP2-SmBiT, MRAP2-LgBiT + MRAP2-SmBiT. PRKACA is a non-interacting cytosolic protein that is used as a negative control. Surprisingly, when MRAP2-LgBiT and MRAP2-SmBiT are transfected together, we see a NanoBiT luminescence signal that is significantly higher than the negative control (Figure 4A). These results show that the C-terminal domains are of MRAP2 are in close proximity to each other in live cells (Figure 4B). MRAP2 Δ38-44, TM-CTD, NTD-TM, TM, and G48L+G52L with C-terminal LgBiT and C-terminal SmBiT also resulted in a significant NanoBiT luminescence signal when compared to the negative control and these results are consistent with the co-immunoprecipitation experiments. Unexpectedly, when LgBiT-MRAP2 and MRAP2-SmBiT are transfected together, there is less NanoBiT signal than the negative control. The absence of a significant NanoBiT signal from the anti-parallel orientation may be due to the inherent size difference between the N- and C-terminal domains of MRAP2, as the C-terminal domain is significantly larger than the N-terminal domain. Once MRAP2 multimers are positioned in the membrane, the shorter length of the N-terminal domain may not favorably allow for a LgBiT and SmBiT interaction. Overall, these results indicate that the C-terminal domains of MRAP2 are in close-proximity in the cell but does not exclude anti-parallel MRAP2 dimers.

**Figure 4.**
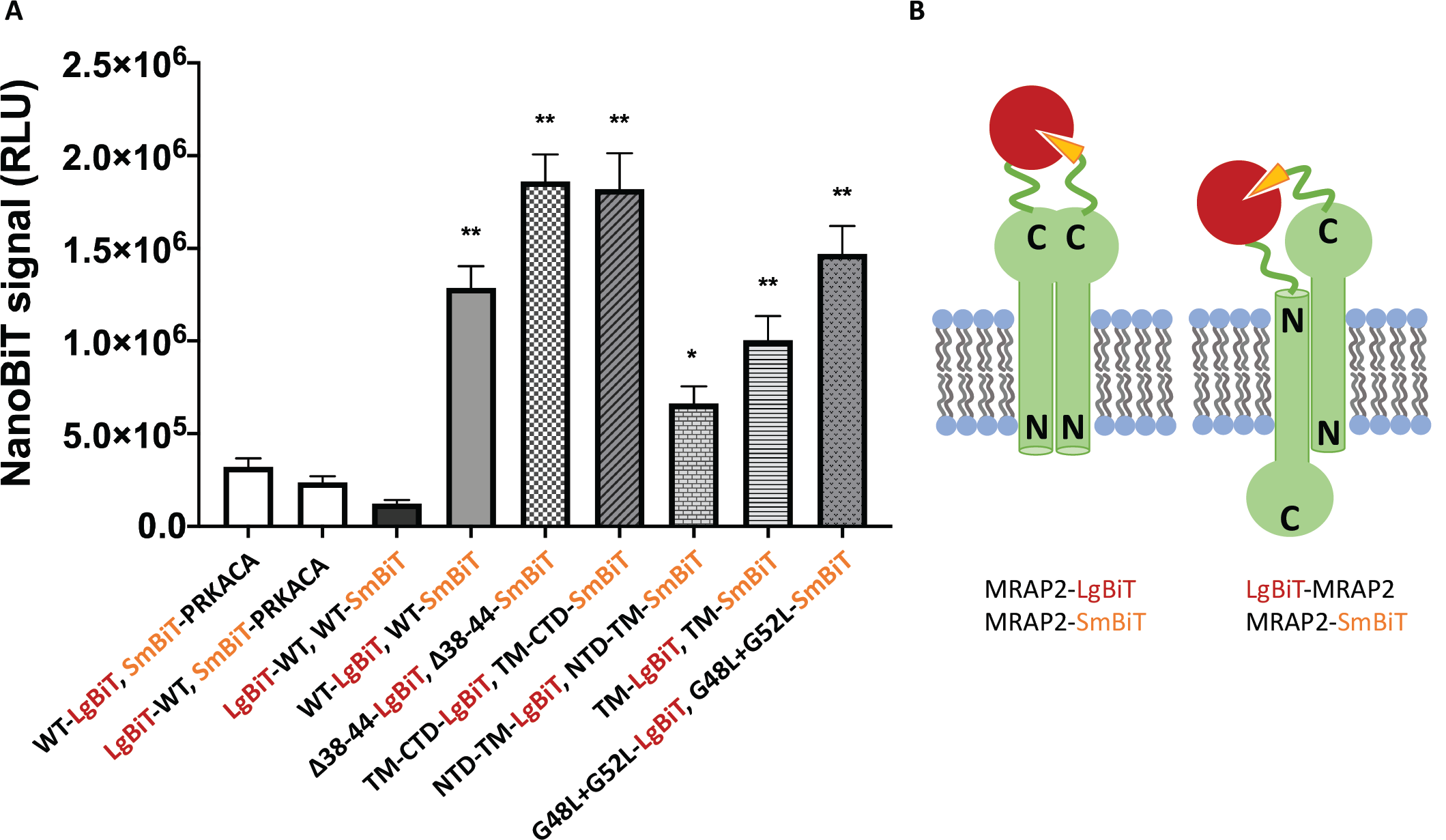
MRAP2 forms parallel dimers or higher order oligomers. A, NanoBiT signal from live cells co-expressing MRAP2-SmBiT with LgBiT-MRAP2 or MRAP2-SmBiT with MRAP-LgBiT. All mutants have both SmBiT and LgBiT as C-terminal fusions. SmBiT-PRKACA is a cytosolic control construct with SmBiT fused to cAMP-dependent protein kinase catalytic subunit alpha. B, Schematic showing reconstituted NanoLuc luciferase from MRAP2 parallel dimers (left) and anti-parallel dimers (right). Bars represent the mean from at least three independent experiments performed in triplicate. Error bars are SEM. Statistical significance of differences was analyzed by *t* test. **p < 0.0001, *p < 0.001 versus WT-LgBiT, SmBiT-PRKACA

To determine whether MRAP2 forms higher order oligomers, whole-cell cross-linking and blue native polyacrylamide gel electrophoresis were employed. Whole-cells expressing FLAG-MRAP2 were cross-linked with a membrane-permeable, lysine-reactive cross-linker and the resulting whole cell lysates were analyzed by SDS-PAGE and western blotting (Figure 5A). The presence of higher molecular weight bands with the apparent molecular weights of cross-linked dimeric and trimeric MRAP2 suggests that MRAP2 forms higher order oligomers that can be cross-linked in whole cells. MRAP2 containing lysates from non-cross-linked cells were also separated on a polyacrylamide gel under native conditions with varying concentrations of *n*-dodecyl-ß-maltoside detergent (Figure 5B). Consistent with the cross-linking results, non-cross-linked MRAP2 forms higher molecular weight structures. These structures are most likely dimers, tetramers, and octamers based on the molecular weights of the bands. Since native protein shape and the amount of bound detergent will affect protein complex mobility through the gel matrix, molecular weights predicted from native gels are approximate. Additionally, the absence of higher order oligomer bands such as tetramers and octamers in the cross-linking experiment are likely a result of incomplete cross-linking of higher order structures. In summary, these results indicate that MRAP2 forms parallel dimers and higher order oligomers such that the C-terminal domains of MRAP2 proteins are in close proximity in the membrane (Figure 5C).

**Figure 5.**
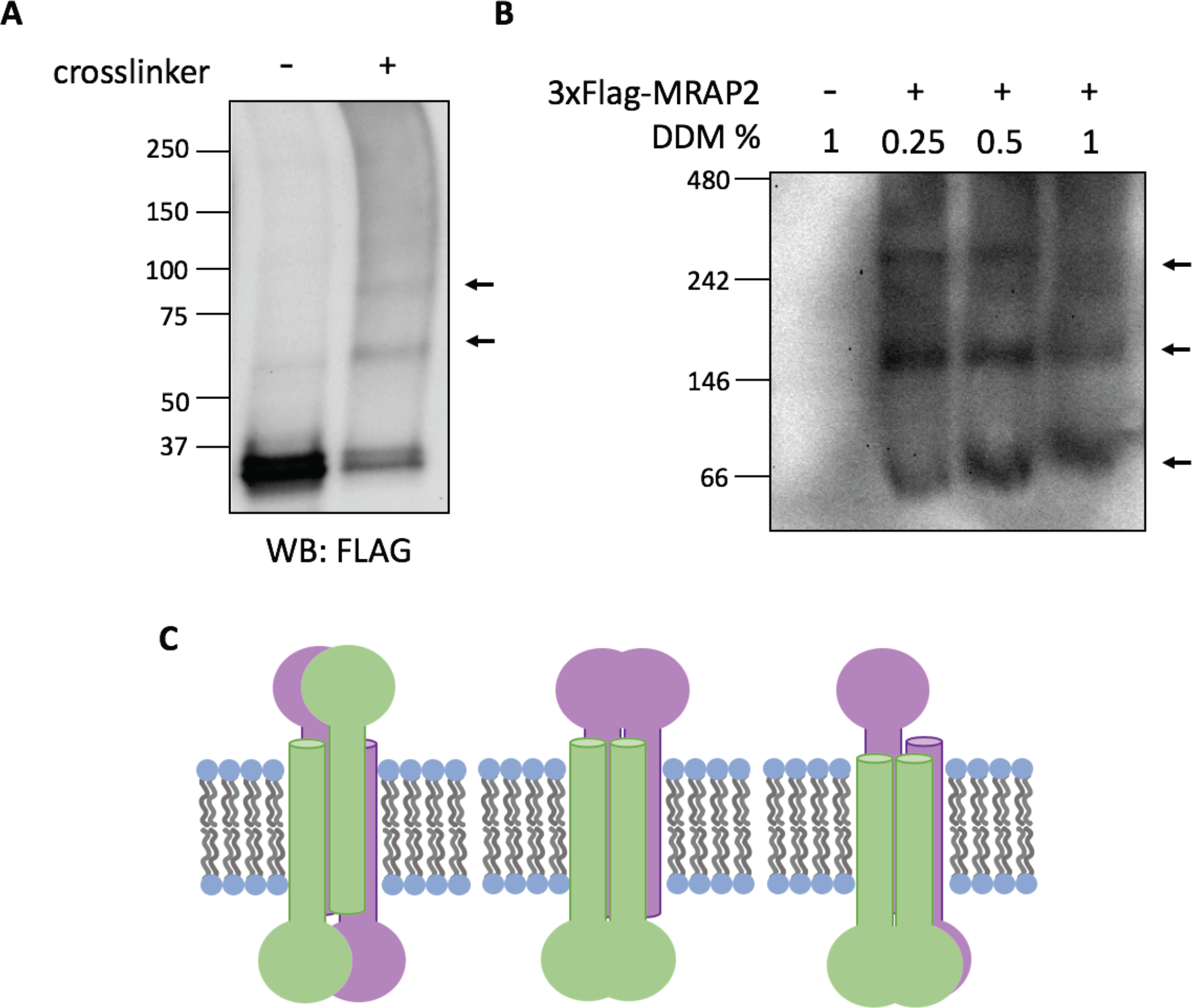
MRAP2 forms higher order oligomers. A, HEK293T cells transfected with FLAG-MRAP2 were incubated with an irreversible cross-linker (DSG or DSS). Proteins in cell lysate were separated by SDS-PAGE and immunoblotted. B, HEK293T cells expressing FLAG-MRAP2 were solubilized with varying concentrations of *n*-dodecyl-ß-maltoside (DDM) and analyzed by blue native PAGE and immunoblotted. The band around 66 kDa is the approximately the molecular weight of a FLAG-MRAP2 dimer. C, Schematic showing possible MRAP2 higher order oligomers.

## DISCUSSION

The N-terminal and transmembrane domains of MRAP2 and MRAP1 are highly conserved and the N-terminal domain of MRAP1 is essential for its dual topology and homodimerization. However, we show that the mechanism that regulates MRAP2 dimerization is distinct from that of MRAP1. Deletion of the conserved polybasic motif that dictates MRAP1’s membrane orientation from MRAP2 does not abolish dual membrane orientation nor dimerization of MRAP2. The “positive-inside” rule that predicts the orientation of transmembrane domains is based on approximately 15 residues on either side of the transmembrane domain and establishes that the more positively-charged side will end up on the cytosolic side of the plasma membrane ^18,19^. In fact, single point mutations in dual topology membrane proteins from bacteria that have a near zero charge bias can shift the orientations of these proteins^23^. Previous experiments that investigated MRAP1’s membrane topology are consistent with both the positive-inside rule and algorithms that predict transmembrane helices and membrane topology such as TMHMM^24^ (Figure S3). Around half of MRAP1 is glycosylated, supporting the fact that MRAP1 can be inserted into the membrane in both N_exo_/C_cyto_ and N_cyto_/C_exo_ orientations. MRAP1’s dual topology has also been validated by bimolecular fluorescence complementation experiments and the presence of both N- and C-terminal antibody epitopes on the cell surface ^15^. Based on the positive-inside rule and computational predictions, MRAP2 is predicted to favor a N_cyto_/C_exo_ orientation. However, this study, along with previous studies^16^ support MRAP2 having dual topology, contrary to the predicted N_cyto_/C_exo_ orientation. MRAP2’s dual topology defies the positive-inside rule. Our study reveals that the molecular features that dictate MRAP1 and MRAP2 orientations are distinctly different. These differences between MRAP1 and MRAP2 highlight the importance of identifying the mechanism behind MRAP2’s dual topology and this will be the subject of future studies.

MRAP2 has been shown to co-immunoprecipitate with several GPCRs including all five melanocortin receptors, the orexin receptor 1 (OX1R), the prokineticin receptor 1 (PKR1), and the growth hormone secretagogue receptor 1a (GHSR1a)^9–12,17^. MRAP2’s role in modulating GHSR1a has now been thoroughly investigated by Rouault and colleagues^12^. They show that MRAP2 alters GHSR1a signaling by inhibiting constitutive activity of the receptor, enhancing ghrelin-mediated G protein signaling, and inhibiting ghrelin-stimulated recruitment of ß-arrestin to GHSR1a. In addition to demonstrating that MRAP2 can bias the signaling of GHSR1a, the regions of MRAP2 responsible for these effects were also elucidated. Specifically, residues 34 to 43 of the N-terminal domain, the transmembrane domain, and the C-terminal domain of MRAP2 are required for potentiation of GHSR1a while residues 24 to 33 and the C-terminal domain are important for inhibition of ß-arrestin recruitment. These results highlight independent mechanisms of receptor-modulation since distinct regions of MRAP2 are required for the enhancement of G protein signaling versus the inhibition of ß-arrestin recruitment. The C-terminal domain of MRAP2 is highly conserved and is necessary for modulation of OX1R, PKR1, and GHSR1a^10,12^. While there is sufficient evidence pointing towards the C-terminus of MRAP2 as being the most important domain for the regulation of GPCRs, until this point, the regions of MRAP2 that are essential for its homodimerization have not been identified. In this study, we show that MRAP2 dimerizes through its transmembrane domain. Interestingly, the transmembrane domain of MRAP2 does not appear to be necessary for OX1R and PKR1 inhibition, but has been shown to be play a role in potentiating GHSR1a signaling^10,12^. An obesity-linked mutation within the transmembrane of MRAP2 has also been reported^4^. Additionally, the transmembrane domain of MRAP1 is essential for MC2R trafficking^13,14^. RAMP1 dimers are inhibited by the presence of the calcitonin receptor-like receptor (CRLR) and RAMP1 forms heterodimers with CRLR at a 1:1 ratio, suggesting that RAMP1 interacts with this receptor as a monomer^25^. A series of MRAP-MC2R or MRAP-MRAP-MC2R fusion proteins were used by Malik et al. to show that MRAP1 dimers were required for MC2R activity^26^. It will be interesting to see if MRAP2 dimerization is required for function in the same manner as MRAP1 or if MRAP2 binds to GPCRs as a monomer like RAMP1. Since MRAP2 is somewhat promiscuous and appears to interact with several GPCRs, it is possible that the oligomeric state of functional MRAP2 is GPCR-specific. Further experimentation is needed to determine whether MRAP2 dimerization is necessary for its modulation of GPCRs, as well as to identify motifs or residues within the transmembrane domain that facilitate dimerization. Transmembrane helix-packing motifs that consist of small residues occurring every four or seven residues have been identified^21,22,27^. EmrE, a small bacterial multidrug transporter, forms antiparallel dimers through conserved glycine residues^28^. However, we find that the glycine residues within MRAP2’s transmembrane domain are not required for dimerization.

Bimolecular fluorescence complementation experiments that incorporate yellow fluorescent protein (YFP) fragments on the N- and C-terminal ends of MRAP1 show that MRAP1 forms exclusively anti-parallel dimers since YFP can only be reconstituted when fragments are on opposite ends and there is no YFP complementation when fragments are place on the same ends of MRAP1^13,14,16^. Similar experiments have been performed for MRAP2 supporting anti-parallel MRAP2 dimers. However, there have been no experiments ruling out a parallel orientation for MRAP2 dimers. Surprisingly, we find that the C-terminal domains of MRAP2 are in close proximity in live cells. This suggests that MRAP2 forms parallel dimers or oligomerizes in such a way that brings the C-terminal domains in close proximity. It is important to note that the absence of data supporting an anti-parallel MRAP2 dimer in our experiments does not eliminate the possibility that MRAP2 can form anti-parallel dimers, rather, it is possible that the size difference between the N- and C-terminal domains of MRAP2 prevents the stable reconstitution of the enzyme used in our complementation experiment once MRAP2 is positioned in the membrane.

We also present evidence for higher-order MRAP2 oligomers that may also explain the presence of parallel MRAP2 dimers. At this juncture, it is not clear whether these higher order oligomers are required for function and it is possible that the oligomeric state of MRAP2 is dependent on the local environment that it exists in. MRAP2 has been shown to form dimers that are resistant to reducing and denaturing conditions from mouse tissue immunoblots^17^. Similarly, the proteolipid protein (PLP), an abundant CNS myelin protein important for the stabilization of myelin membranes, also forms SDS-resistant dimers in the ER^29^. PLP forms higher order oligomers only after reaching the cell surface. An attractive hypothesis is that MRAP2 facilitates the trafficking of GPCRs to the cell surface and once at the cell surface, the changes in MRAP2’s oligomeric state could act as a molecular switch to tune its regulation over GPCRs.

Currently, there is very little information regarding MRAP2 structure and it is not uncommon for single-pass transmembrane proteins to have intrinsically disordered domains making them difficult to study using classic biophysical techniques ^30^. These disordered regions often have functional importance since this flexibility allows them to interact with multiple protein partners. While the transmembrane domain of MRAP2 is most likely a transmembrane helix, there is very little predicted secondary structure in the N- and C-terminal domains. Since MRAP2 has been shown to interact with several receptors, it is possible that in the presence of these receptors, MRAP2 adopts a more rigid conformation allowing high resolution structures to become more obtainable.

MRAP2 and MRAP1 are the only known eukaryotic proteins that adopt a highly unique dual topology in the membrane. While the N-termini and transmembrane domains of these homologs are highly conserved, we show key differences between MRAP2 and MRAP1 membrane orientation and oligomerization. Specifically, the conserved polybasic motif that is essential for MRAP1’s anti-parallel orientation and dimerization does not dictate the topology and oligomeric state of MRAP2. Additionally, for the first time, we provide evidence for the transmembrane domain as being the minimal dimerization domain and identify a new parallel orientation for MRAP2 oligomers. Elucidating the molecular framework behind MRAP2 structure will give insight into the mechanisms by which other single-pass transmembrane proteins and accessory proteins modulate their receptors. Furthermore, given the essential role of MRAP2 in the modulation of GPCRs that are critical for the maintenance of energy homeostasis, understanding the structure of MRAP2 will aid in unraveling the complex neural circuitry responsible for the central regulation of energy balance.

## EXPERIMENTAL PROCEDURES

### Expression Constructs

3xFlag-tagged wild-type MRAP2 constructs are in pSF vectors and 3xHA-tagged RAMP3 expression constructs are in pcDNA3.1 vectors. Wild-type expression constructs were kindly provided by Dr. Roger Cone (University of Michigan, Life Science Institute, Ann Arbor, MI). Mutations in MRAP2 constructs were generated using PCR primer-based site-directed mutagenesis, using primers generated manually and by www.primerdesigner.com^31^. NanoBit vectors were purchased from Promega (Madison, WI). MRAP2 was cloned into NanoBiT vectors by directional cloning with the following restriction sites: SacI and XhoI for C-terminal LgBiT and SmBiT, SacI and NheI for N-terminal LgBiT and SmBiT. All constructs were verified by DNA sequencing.

### Cell culture and transfections

HEK293T cells (ATCC CRL-3216, Lot # 62729596) were cultured in Dulbecco’s Modified Eagles Medium (DMEM) and CHO cells were cultured in F-12 medium. Both cell lines were cultured in media supplemented with 10% fetal bovine serum and GlutaMAX from Life Technologies (Carlsbad, CA) at 37 °C in 5% CO_2_. Cells were transfected with Lipofectamine 2000 DNA transfection reagent from Invitrogen (Carlsbad, CA) 18-24 hours after plating. All experiments were performed 24 hours post-transfection.

### Co-immunoprecipitation and western blot

HEK293T cells were co-transfected with N-terminally HA-tagged and N-terminally Flag-tagged MRAP2 wild-type or mutant constructs using Lipofectamine 2000 following manufacturer’s instructions. After 24 hours post-transfection, cells were washed with ice-cold phosphate buffered saline (PBS, pH 7.4) and lysed with 0.2% *n*-dodecyl-ß-maltoside (DDM) in PBS with EDTA and HALT protease inhibitors (Pierce Biotechnology, Rockford, IL) added. The lysed cells were incubated on ice for 30 minutes and the lysate was clarified at 17,000 x g at 4°C for 25 minutes. Protein concentrations were determined using a BCA protein quantitation kit (Pierce Biotechnology, Rockford, IL) according to the manufacturer’s instructions. Samples treated with PNGase F (New England Biolabs, Ipswich, MA) were treated according to the manufacturer’s instructions under denaturing reaction conditions. For immunoprecipitations, lysates were incubated with either anti-FLAG M2 agarose beads (Sigma-Aldrich, St. Louis, MO) or anti-HA HA-7 agarose beads (Sigma-Aldrich, St. Louis, MO) overnight at 4°C with end-over-end mixing. The same volume of lysate was used for anti-Flag and anti-HA pull-downs. The beads were washed four times with 0.1% % DDM in PBS. The protein from the beads were either eluted with Laemmli buffer with DTT and boiled for 5 mins or eluted with Laemmli buffer without DTT and reduced with DTT for 30 minutes at 37 °C after separating the beads from the sample. The eluted proteins or whole cell lysates were separated by SDS-polyacrylamide gel electrophoresis on a 4-20% gradient gel (Bio-Rad Laboratories, Hercules, CA) and transferred to polyvinylidene fluoride (PVDF) membranes. The membranes were blocked overnight at 4°C with 5% bovine serum albumin (BSA) in tris-buffered saline with 0.1% Tween-20 (TBST). Either mouse M2 Flag or mouse HA-7 primary antibodies (Sigma-Aldrich, St. Louis, MO) were used at a 1:1000 dilution in 2.5% BSA in TBST for 1 hour at room temperature. The blots were washed (4X, 3 min) with TBST. A secondary goat anti-mouse HRP (horse radish peroxidase)-conjugated antibody (Abcam, Cambridge, United Kingdom) was used at a 1:10,000 dilution for 1 hour at room temperature. The blots were washed (4X, 3 min) with TBST before adding the ECL western blotting substrate (Pierce Biotechnology, Rockford, IL). The blots were imaged using a ChemiDoc™ XRS+ System and analyzed using Image Lab™ Software (Bio-Rad Laboratories, Hercules, CA).

### Immunostaining and confocal microscopy

HEK293T cells were plated in 8-well chamber slides (ibidi GmbH, Martinsreid, Planegg, Germany) previously coated with poly-D-lysine. Cells were transfected with either N-terminally Flag-tagged MRAP2 or MRAP2 Δ38-44 and C-terminally Flag-tagged MRAP2 or MRAP2 Δ38-44. After 24 hours post-transfection, transfected cells were washed twice with ice-cold PBS and fixed with 2% formaldehyde in PBS for 10 minutes at room temperature. Cells were washed with PBS (3X, 5 min) and permeabilized samples were incubated with 0.5% saponin in PBS for 10 minutes. Unpermeabilized cells were incubated with PBS for 10 minutes. Cells were washed with PBS (3X, 5 min). Cells were then blocked with 1% BSA in PBS for unpermeabilized cells or 1% BSA, 0.1% saponin in PBS for permeabilized cells for 30 minutes. Blocked cells were then incubated with a mouse M2 Flag antibody (Sigma-Aldrich, St. Louis, MO) at a 1:600 dilution in 1% BSA in PBS for 1 hour. Cells were washed again with PBS (3X, 5 min) and incubated with a goat anti-mouse Alexa 568 antibody (highly cross adsorbed) (Invitrogen, Carlsbad, CA) at a 1:1000 dilution in 1% BSA in PBS for 1 hr in the dark. For the fixation control experiments, cells were incubated with an alpha tubulin monoclonal antibody conjugated to Alexa Fluor 488 (B-5-1-2, Invitrogen, Carlsbad, CA) at 2 *μ*g/mL in 1% BSA in PBS for 1 hour. Cell were washed with PBS (3X, 5 min), stained with Hoechst 33342 (5 ug/mL) (Invitrogen, Carlsbad, CA) for 1 min, washed with PBS again (3X, 5 min) and slides were mounted with Fluoromount-G mounting medium (SouthernBiotech, Birmingham, AL). Images were acquired on a Leica SP5 confocal microscope and analyzed by Fiji ImageJ.

### Immunostaining and flow cytometry

HEK293T cells were transfected with either N-terminally Flag-tagged MRAP2 or MRAP2 Δ38-44, C-terminally Flag-tagged MRAP2 or MRAP2 Δ38-44, N-terminally HA-tagged RAMP3, or C-terminally HA-tagged RAMP3. After 24 hours post-transfection, transfected cells were washed twice with ice-cold PBS and fixed with 2% formaldehyde in PBS for 15 minutes at room temperature. Fixed cells were then washed twice with PBS and split into two separate samples. Each of the samples was further washed with either FACS buffer without Triton X-100 (PBS pH 7.4 with 1% BSA, 1 mM EDTA, 0.05% sodium azide) for non-permeabilized samples or FACS buffer with Triton X-100 (PBS pH 7.4 with 1% BSA, 1 mM EDTA, 0.05% sodium azide, 0.1% Triton X-100) for permeabilized samples. Unpermeabilized cells were resuspended in FACS buffer and permeabilized cells were resuspended in FACS buffer with Triton X-100. Alexa Fluor 488 conjugated anti-HA (Cell Signaling Technologies #2350, Danvers, MA) or anti-FLAG (Cell Signaling Technologies #15008, Danvers, MA) antibodies were added at a 1:50 dilution and cell were incubated with the fluorescent antibodies for 1 hour in the dark at 4°C. Stained cells were then washed with FACS buffer three times and resuspended in FACS buffer before flow analysis. Cells were analyzed on a BD LSRII and 10,000 cells were counted and analyzed for each sample. FlowJo was used for data processing. The median fluorescence intensity (MFI) value for the unpermeabilized cells was divided by the MFI of the permeabilized samples to normalize for differences in protein expression levels.

### NanoBiT protein-protein interaction assay

HEK293T cells were plated in an opaque, white 96-well plate. Cells were transfected with Lipofectamine 2000 according to manufacturer’s instructions. After 24 hours post-transfection, media is removed and replaced with Opti-MEM (Life Technologies, Carlsbad, CA). NanoBiT live-cell substrate was added according to manufacturer’s instructions and incubated for 5 minutes. Luminescence was measured using a Perkin-Elmer EnVision plate reader.

### Whole-cell cross-linking

HEK293T cells were transfected with 3xFlag-MRAP2. After 24 hours post-transfection, cells were washed with PBS and incubated with 1 mM disuccinimidyl suberate or disuccinimidyl glutarate (Thermo Fisher) in PBS for 30 minutes at room temperature. The cross-linking reaction was quenched with 30 mM Tris-HCl in PBS for 10 minutes. Cells were then pelleted and washed with 30 mM Tris-HCl in PBS. The washed cell pellet was lysed with 1% DDM in PBS and analyzed by SDS-PAGE and western blotting as described in the *co-immunoprecipitation and western blot* section.

### Native protein gel electrophoresis and western blot

HEK293T cells were transfected with 3xFlag-MRAP2. At 24 hours post-transfection, cells were washed with PBS and lysed with a native lysis buffer (50 mM BisTris pH 7.2, 50 mM NaCl, 10% glycerol) with protease inhibitors and varying concentrations of DDM for 30 minutes on ice. Lysates were clarified at 17,000 x g at 4°C for 25 minutes. Protein concentrations were determine using a BCA protein quantitation kit (Pierce Biotechnology, Rockford, IL) according to the manufacturer’s instructions. A 10x loading dye (5% w/v Coomassie blue G-250, 500 mM 6-aminohexanoic acid) was added to lysates right before loading samples into a NativePAGE Bis-Tris gel 4-16% (Life Technologies). The proteins were separated according to Wittig et al.^32^ and transferred onto a PVDF membrane using a standard Tris-glycine transfer buffer with 0.05% SDS. The membrane was de-stained in methanol for 3 minutes. The membrane was blocked and immunoblotted as described in the *co-immunoprecipitation and western blot* section.

## DATA AVAILABILITY

All data for this study are available within the article.

## ACKNOWLEDGEMENTS

We thank Professor Julien A. Sebag for his advice and expertise regarding MRAP2, Ben Abrams for his microscopy support, and Bari H. Nazario for her help with flow cytometry. This project was funded by NIH grants R01DK110403 to Glenn L. Millhauser and R01DK070332 to Roger D. Cone.

