## Supplemental Figures 1-3 for "Membrane Orientation and Oligomerization of the Melanocortin Receptor Accessory Protein 2"

**Supporting Information**

**Figure S1**

**Figure S2**

**Figure S3**

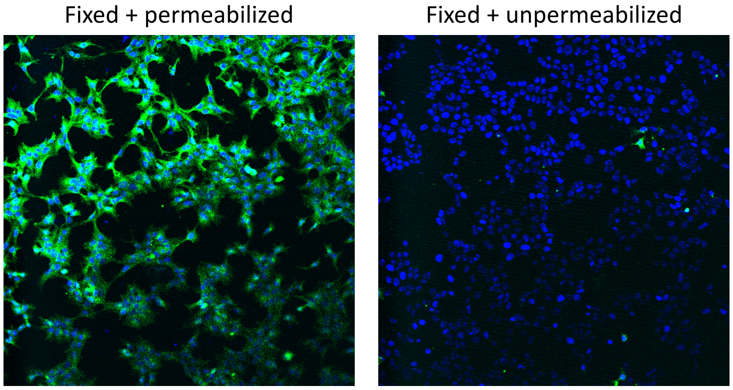


**Supplemental Figure 1 Cell fixation with formaldehyde results in minimal membrane permeabilization.** HEK293T cells were fixed with 2% formaldehyde and incubated with an antibody against alpha-tubulin conjugated to Alexa Fluor 488 and Hoechst 33342, a membrane permeable nuclear stain. Hoechst 33342 allows for nuclear staining of both permeabilized and unpermeabilized cells. Intracellular alpha-tubulin is shown in green and the nuclear DNA is shown in blue. The unpermeabilized cells (right) show negligible staining of intracellular alpha-tubulin as compared to the permeabilized cells (left).

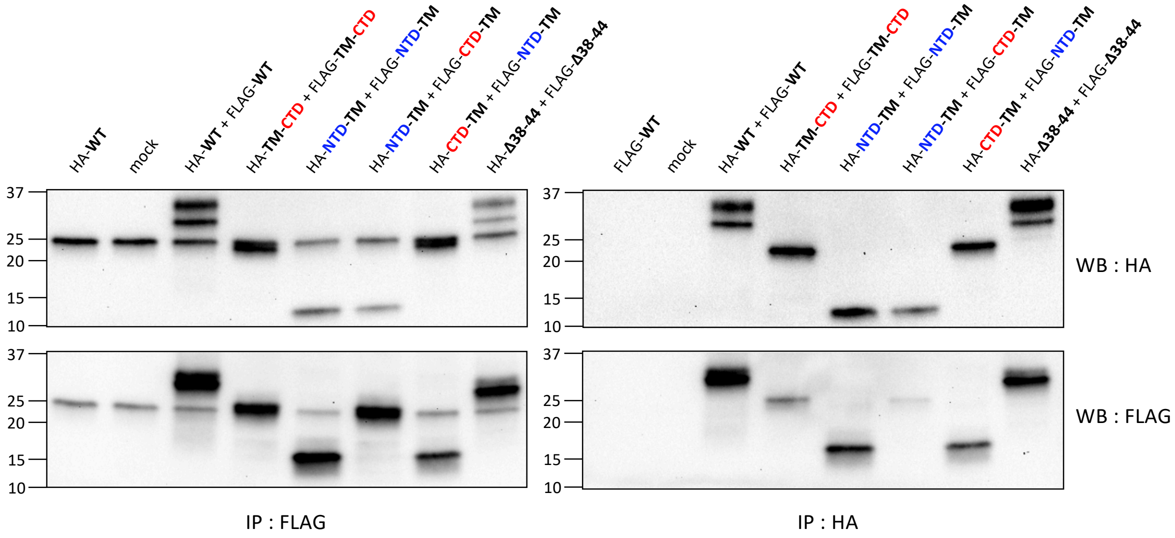


**Figure S2 Co-immunoprecipitation of MRAP2 HA- and FLAG-tagged constructs from CHO cells.** CHO cells are co-transfected with HA- and FLAG-tagged MRAP2 constructs. Co-immunoprecipitations from cell lysates followed by immunoblotting show that neither the N-terminal domain, nor the C-terminal domain, nor residues 38 to 44 are required for MRAP2 dimerization.

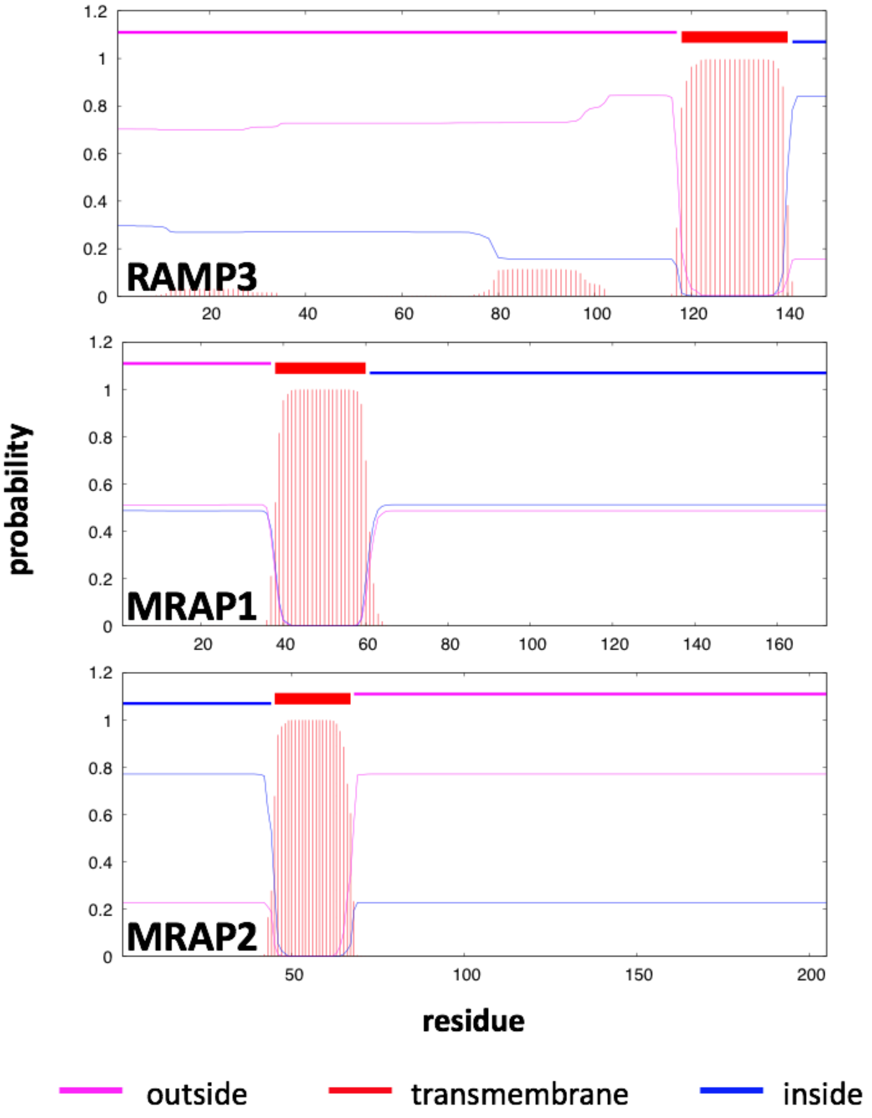


**Figure S3 MRAP2 is predicted to favor an $N_{cyto}/C_{exo}$ orientation.** The transmembrane helix and topology for RAMP3, MRAP1, and MRAP2 were predicted using TMHMM2.0.
